## Supplementary material for "A circular zone of attachment to the extracellular matrix provides directionality to the motility of *Toxoplasma gondii* in 3D": Suppl. Figures 1-9

Suppl. Figure 1

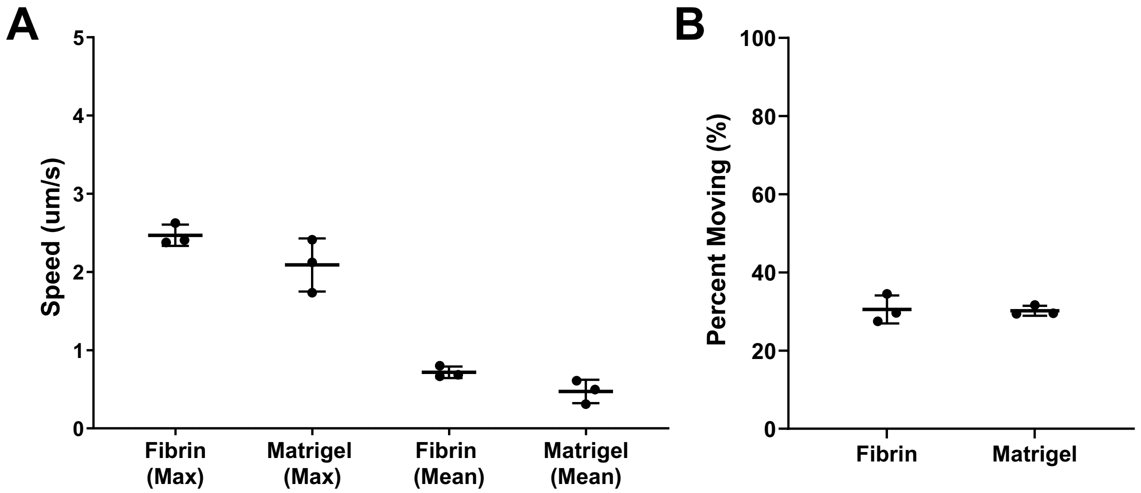

Suppl. Figure 2

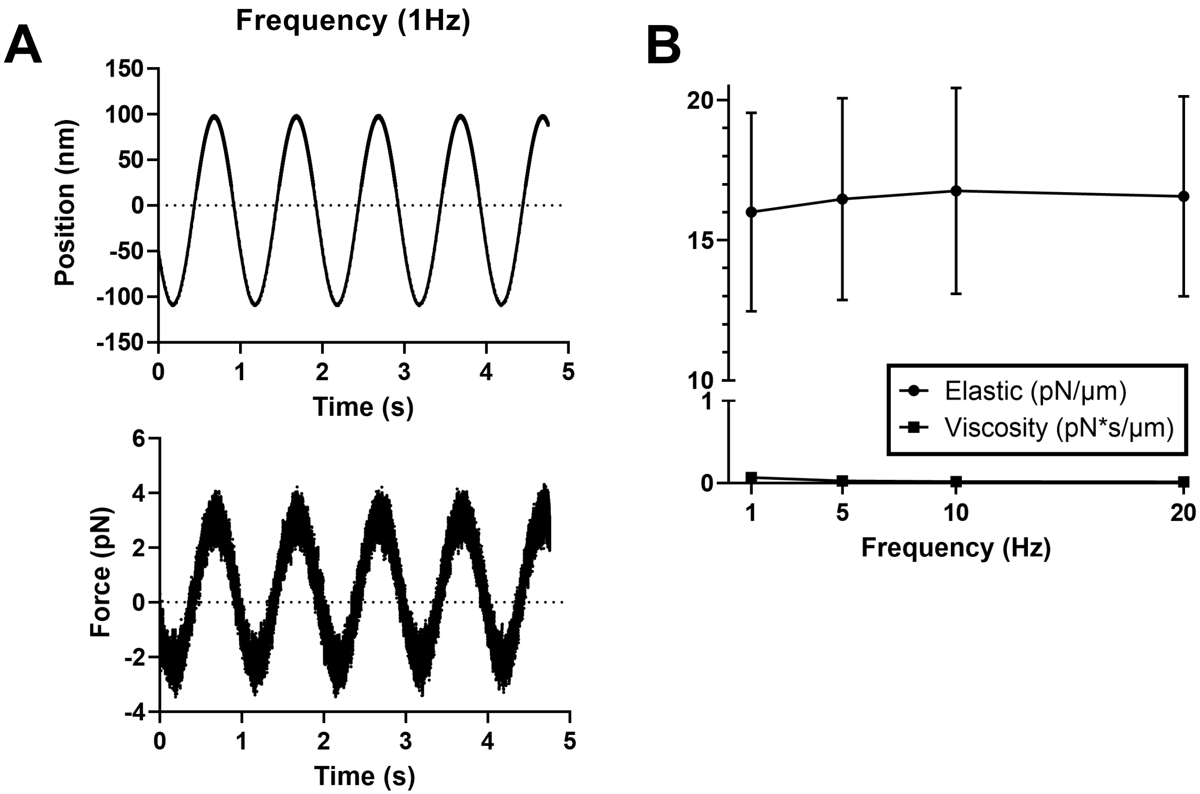

Suppl. Figure 3

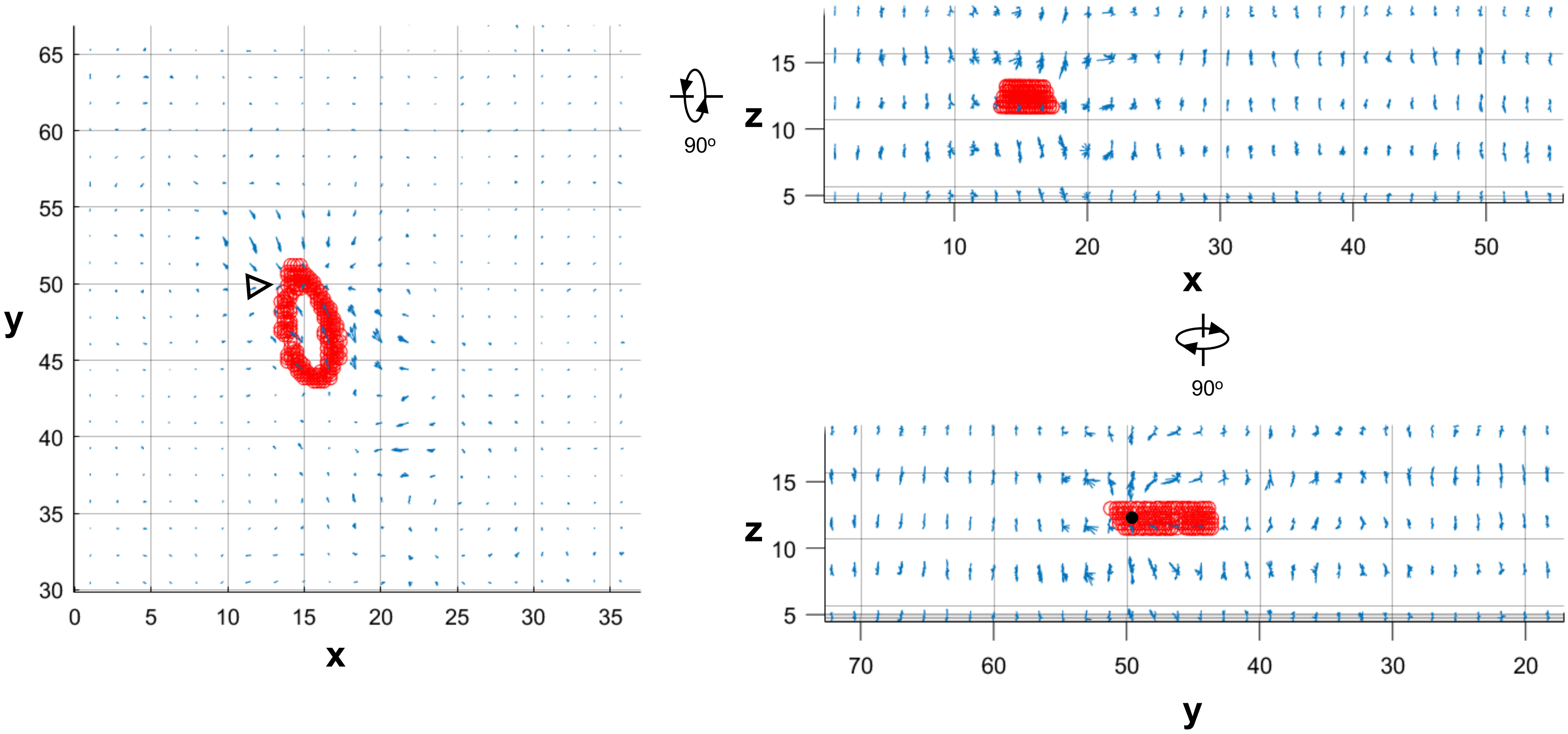

Suppl. Figure 4

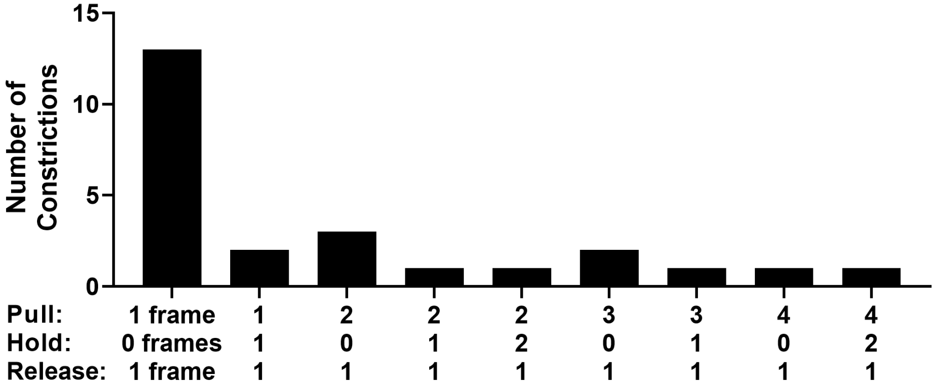

### Suppl. Figure 5

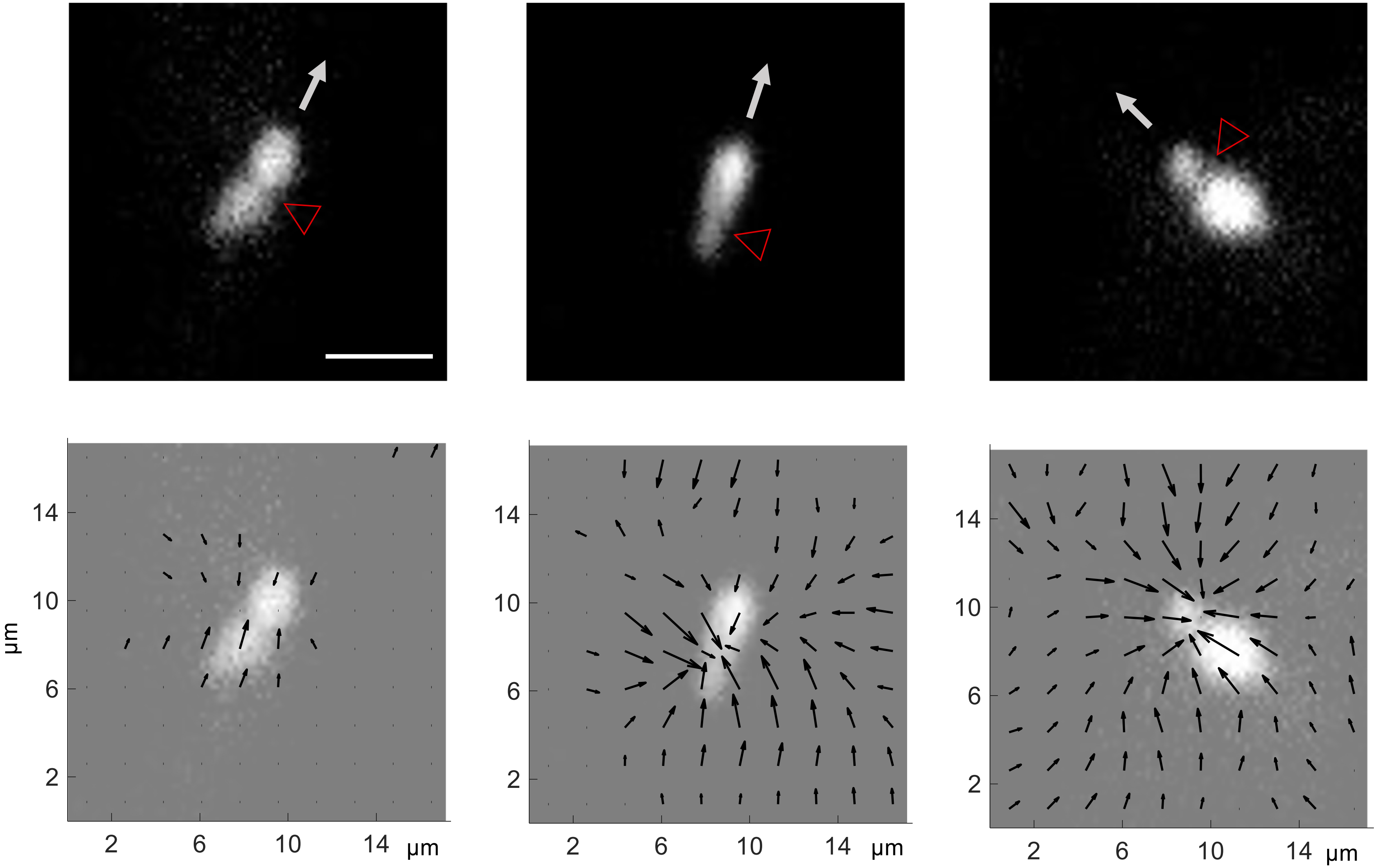

Suppl. Figure 6

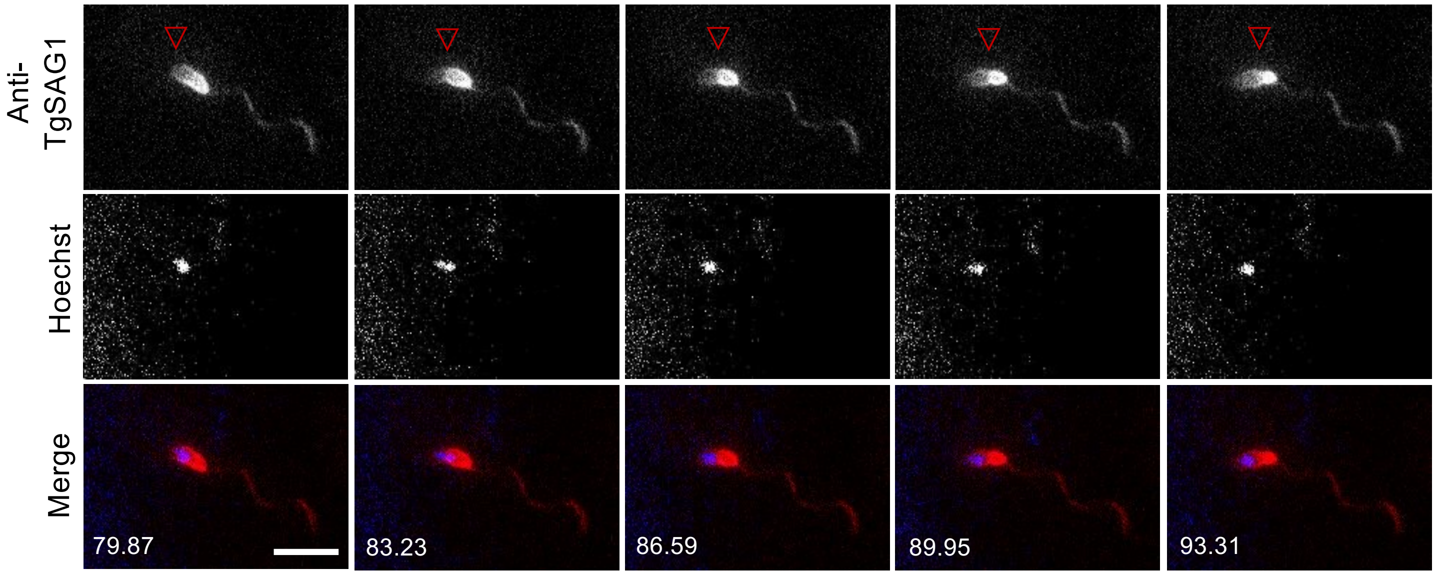

Suppl. Figure 7

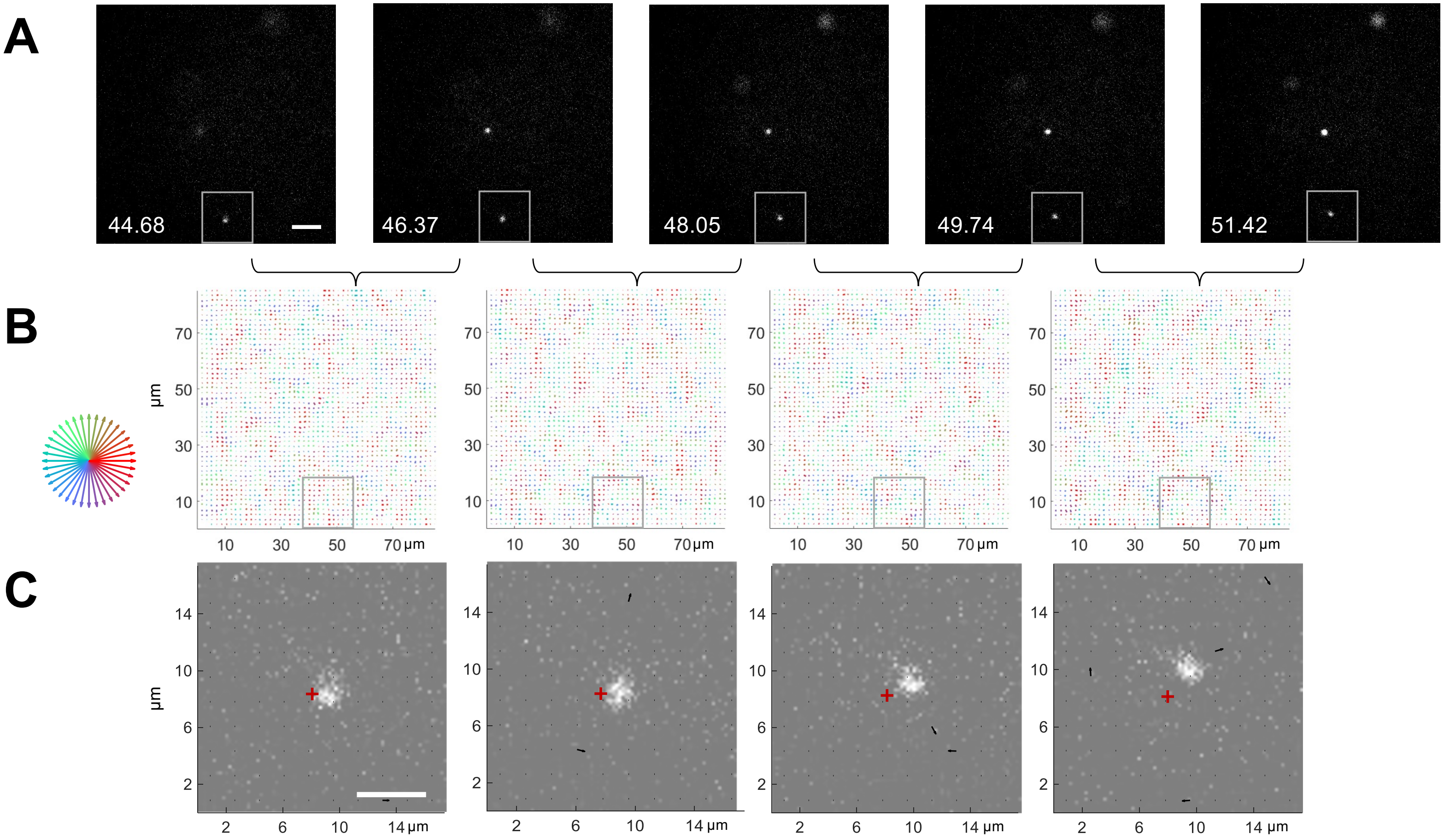

Suppl. Figure 8

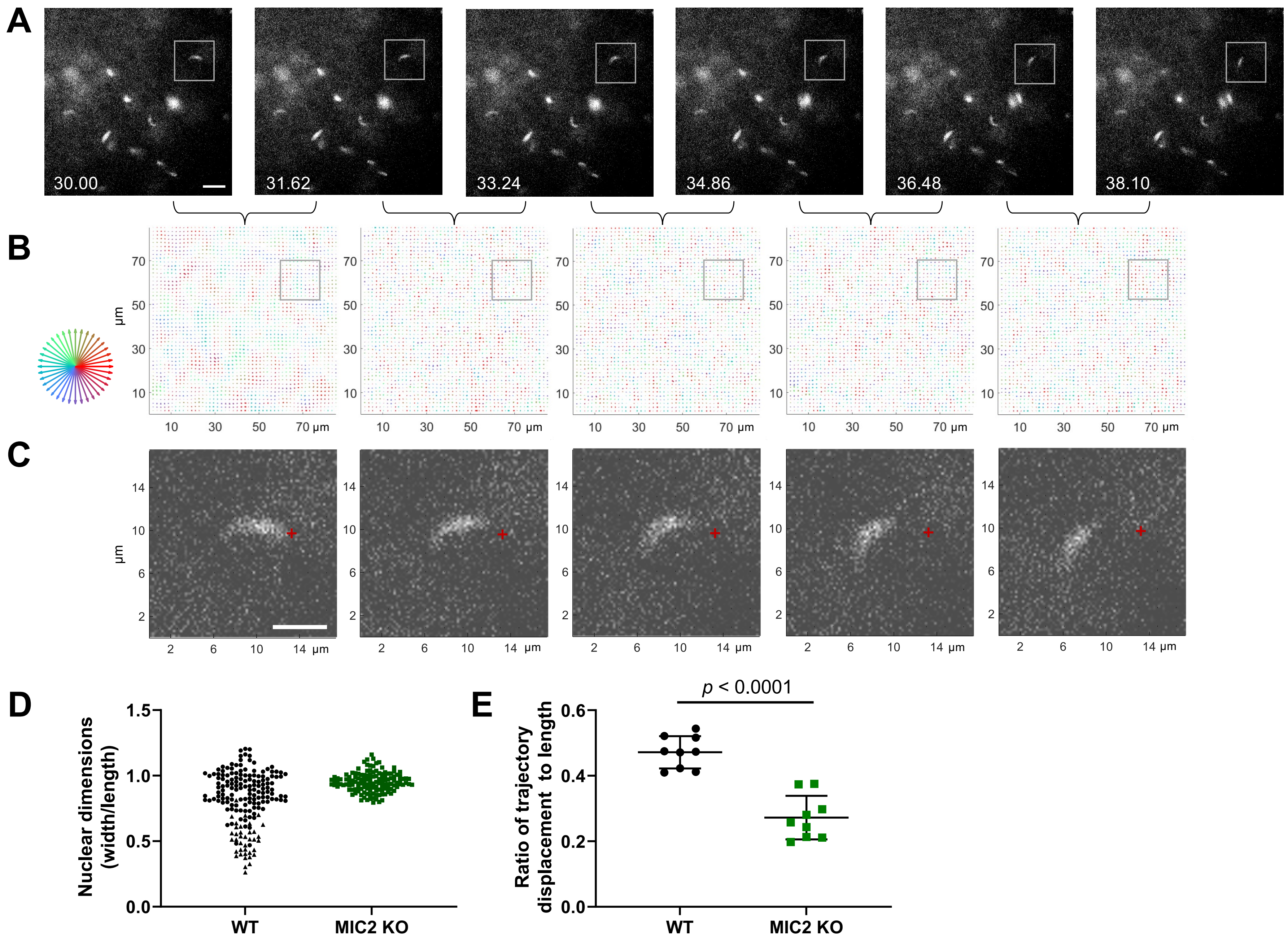

Suppl. Figure 9

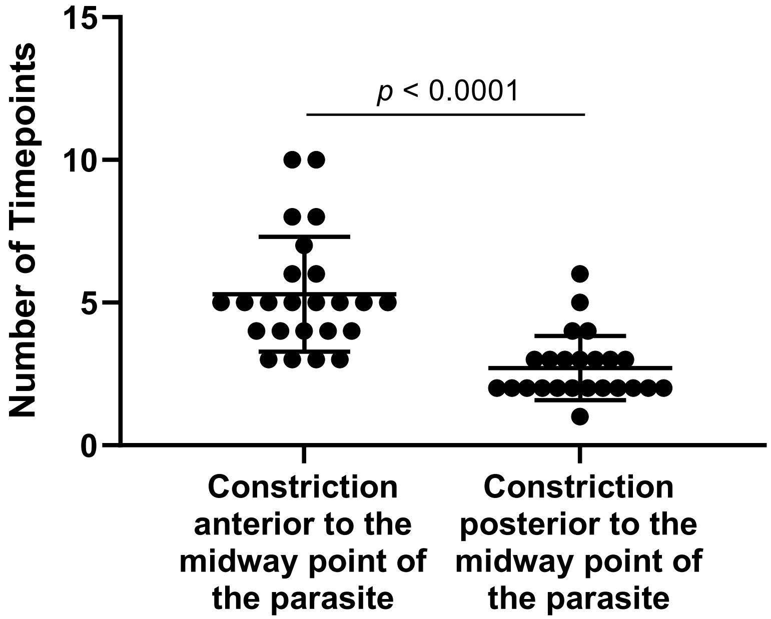
